# Synergistic activity of dispersin B and benzoyl peroxide against *Cutibacterium acnes* biofilms

**DOI:** 10.1101/2023.12.14.571714

**Authors:** Jeffrey B. Kaplan

## Abstract

*Cutibacterium acnes* has been implicated in the pathogenesis of acne vulgaris. *C. acnes* forms biofilms which may contribute to host colonization and antimicrobial resistance. Poly-*N*-acetylglucosamine (PNAG) is an exopolysaccharide that mediates *C. acnes* biofilm formation. In this study we investigated the ability of the PNAG-degrading enzyme dispersin B to sensitize *C. acnes* biofilms to killing by the anti-acne agent benzoyl peroxide (BPO). *C. acnes* biofilms were cultured aerobically in glass tubes in the presence of *Staphylococcus epidermidis* which has been shown to stimulate *C. acnes* biofilm formation. Biofilms were treated with 5-80 μg/ml dispersin B and/or 0.1-2.5% BPO. Treatment of biofilms with dispersin B or BPO alone resulted in a 1-2 log reduction *C. acnes* CFUs, whereas treatment of biofilms with dispersin B followed by BPO resulted in a >6 log reduction in *C. acnes* CFUs. Concentrations as low as 5 μg/ml dispersin B and 0.5% BPO efficiently eradicated *C. acnes* from the dual-species biofilm. Our findings confirm that PNAG protects *C. acnes* from benzoyl peroxide killing and demonstrate that dispersin B and BPO act synergistically to kill *C. acnes* biofilm cells. Dispersin B may be a useful adjunct to BPO for the treatment and prevention of acne.

## INTRODUCTION

The Gram-positive bacterium *Cutibacterium acnes* was first isolated from a patient with acne vulgaris more than 100 years ago (Achermann *et al*., 2014). *C. acnes* is an aerotolerant anaerobe that utilizes sebum and keratin as nutrient sources, making it well adapted for life inside hair follicles. Although some *C. acnes* cellular components such as porphyrins, surface proteins, cytotoxic exoenzymes, and biofilms have been shown to trigger inflammation in the later stages of acne (Mayslich *et al*., 2021), the precise role of *C. acnes* in the early stages of acne remains unknown. For example, *C. acnes* colonizes hair follicles in all people, and several previous studies found no difference in the abundance of *C. acnes* in subjects with or without acne (Barnard *et al*., 2016; Greydanus *et al*., 2021). Recent studies suggest that acne might be the result of an unbalanced equilibrium between *C. acnes* and *Staphylococcus epidermidis* (Bardard *et al*., 2016; Claudel *et al*., 2019; Dréno *et al*., 2017; Lee *et al*., 2019; McLaughlin *et al*., 2019; O’Neill & Gallo, 2018). In one study that identified bacteria in hair follicles from acne patients and healthy controls using a highly sensitive cyanoacrylate biopsy method (Bek-Thomsen *et al*., 2008), all subjects harbored *C. acnes* within their follicles, but the follicles of acne patients also harbored *S. epidermidis*.

Biofilms are defined as densely packed layers of bacterial cells growing attached to a tissue or surface (Costerton *et al*., 1999). Bacteria in a biofilm are encased in a sticky, self-synthesized, extracellular polymeric matrix that holds the cells together in a mass, attaches them to the underlying surface, and protects them from killing by antimicrobial agents and host immunity. Biofilms play a role in many chronic infections which are often difficult to treat because of the protective nature of biofilms. *In vivo, C. acnes* biofilms have been observed in acne lesions (Jahns *et al*., 2012) and *S. epidermidis* biofilms have been observed on skin and in sweat glands (Allen & Mueller, 2011). Both species have been shown to form biofilms *in vitro* (Achermann *et al*., 2014; Otto, 2009) and on the surfaces of implanted medical devices *in vivo* (Bayston *et al*., 2006; Büttcner *et al*., 2015).

Poly-*N*-acetylglucosamine (PNAG) is an extracellular polysaccharide that mediates biofilm formation, antimicrobial resistance, host colonization, immune evasion, and stress tolerance in a wide range of Gram-negative and Gram-positive bacterial pathogens (Cywes-Bentley *et al*., 2013; Soliman *et al*., 2018). PNAG is an essential virulence factor for *Staphylococcus aureus* in mouse models of systemic infection (Kropec *et al*., 2005); for *Aggregatibacter actinomycetemcomitans* in a rat model of periodontitis (Shanmugam *et al*., 2015); for *Klebsiella pneumoniae* in a mouse model of intestinal colonization and systemic infection (Chen *et al*., 2014); and for *Actinobacillus pleuropneumoniae* during natural porcine pleuro-pneumonia infection in pigs (Subashchandrabose *et al*., 2013). Anti-PNAG antibodies have been shown to protect mice against local and/or systemic infections caused by *Streptococcus pyogenes, Streptococcus pneumoniae, Listeria monocytogenes, Neisseria meningitidis* serogroup B, and *Candida albicans* (Cywes-Bentley *et al*., 2013), suggesting that PNAG is an important virulence factor in these organisms. Previous studies showed that both *C. acnes* and *S. epidermidis* produce PNAG (Kaplan *et al*., 2004; Kaplan, 2023a).

Dispersin B is an enzyme that hydrolyzes PNAG (Kaplan *et al*., 2004). Dispersin B has been shown to inhibit the attachment of *C. acnes* cells to surfaces, to inhibit *C. acnes* biofilm formation, and to sensitize *C. acnes* biofilms to killing by tetracycline and benzoyl peroxide *in vitro* (Kaplan, 2023a). Dispersin B has also been shown to inhibit attachment of *S. epidermidis* cells to pig skin *in vivo* (Kaplan *et al*., 2018), and to inhibit *S. epidermidis* biofilm formation, detach preformed *S. epidermidis* biofilms, and sensitize preformed *S. epidermidis* biofilm cells to killing by cetylpyridinum chloride and rifampicin *in vitro* (Ganeshnarayan *et al*., 2009; Lee *et al*., 2008).

Previous studies showed that *C. acnes* forms significantly more biofilm when co-cultured aerobically with *S. epidermidis* than it does when cultured anaerobically as a mono-species biofilm (Kaplan, 2023b). These aerobic *S. epidermidis*/*C. acnes* dual-species biofilms may emulate many aspects *C. acnes* biofilm formation on the dermis, which is well-oxygenated, and in sebaceous glands and hair follicles, which are moderately to severely hypoxic but not anoxic (Evans *et al*., 2006). Previous studies also showed that dispersin B detaches pre-formed aerobic *S. epidermidis*/*C. acnes* dual-species biofilms from glass tubes (Kaplan, 2023b), suggesting that PNAG is a major adhesin in these dual-species biofilms. In the present study we investigated the ability of dispersin B to sensitize aerobic *S. epidermidis*/*C. acnes* dual-species biofilms to killing by the anti-acne agent benzoyl peroxide.

## MATERIALS AND METHODS

### Bacterial strains

The bacterial strains used in this study were *Cutibacterium acnes* HL086PA1 (Fitz-Gibbon *et al*., 2013) and *Staphylococcus epidermidis* strain 5 (Chaingon *et al*., 2008). *C. acnes* HL086PA1 is an erythromycin-resistant strain isolated from a patient with severe acne. It was obtained through BEI Resources (Manassas VA, USA) as part of the NIAID NIH Human Microbiome Project. *S. epidermidis* strain 5 is an erythromycin susceptible, PNAG-producing strain isolated from an implant infection that was previously shown to facilitate growth of *C. acnes* HL086PA1 biofilms *in vitro* (Kaplan, 2023b). Bacteria were routinely passaged aerobically (*S. epidermidis*) or anaerobically (*C. acnes*) on Tryptic Soy agar (Becton, Dickinson) at 37°C. Anaerobic conditions were created using a BD GasPak EZ Anaerobe sachet system.

### Biofilm culture

Inocula for biofilm cultures were prepared by transferring a 1-μl-size loopful of *S. epidermidis* cells from a 24-h-old agar plate into 200 μl of phosphate buffered saline (PBS), and then transferring a loopful of *C. acnes* cells from 72-h-old agar plate into the same tube. The cells were mixed by vortex agitation, diluted 1:1,000 in filter sterilized Tryptic Soy broth (Becton, Dickinson), and then passed through a 5-μm pore-size syringe filter to remove large clumps of cells. Diluted inocula contained 10^6^-10^7^ CFU/ml of each species. Bacteria were present at a *S. epidermidis*:*C. acnes* ratio of 2-10:1. Filtered inocula were aliquoted into sterile 13 × 100 mm glass tubes (1 ml/tube) and incubated for 24 h statically at 37°C in air.

### Dispersin B

Two formulations of dispersin B were employed. The first was a dispersin B solution which contained 1 mg/ml recombinant dispersin B protein dissolved in 50 mM phosphate buffer (pH 5.8), 100 mM NaCl, and 50% glycerol. Dispersin B solution was diluted in PBS to achieve a working concentration of 80 μg/ml enzyme. The second formulation was a dispersin B hydrogel which contained 80 μg/ml dispersin B in a mixture of non-ionic surfactant (poloxamer 407), glycerol, levulininc acid, anisic acid, and phosphate buffer. The dispersin B hydrogel was used directly or diluted with blank hydrogel (no enzyme) to achieve working concentrations of 40, 20, 10 and 5 μg/ml enzyme. Dispersin B solution, dispersin B hydrogel, and blank hydrogel were provided by Kane Biotech (Winnipeg MB, Canada).

### Benzoyl peroxide

CVS Health brand Maximum Strength Acne Treatment Gel containing 10% benzoyl peroxide (BPO) was employed. The gel was diluted in PBS to obtain final working concentrations of 2.5%, 1%, 0.5%, and 0.1% BPO.

### Two-step biofilm treatment and enumeration of biofilm CFUs

After growth of biofilms for 24 h, the media was gently aspirated using a finely drawn Pasteur pipette. Each tube was rinsed twice with 3 ml of PBS and gently aspirated after each rinse. For the first treatment, tubes were filled with 1 ml of PBS; 80 μg/ml dispersin B solution; dispersin B hydrogel containing 5, 10, 20, 40 or 80 μg/ml dispersin B; or 1% or 2.5% BPO, as indicated. Tubes were incubated for 15 min at 37°C and then rinsed twice with PBS and aspirated as described above. All treatment and rinse steps involving dispersin B hydrogel were carried out on ice to maintain the liquid state of the hydrogel. For the second treatment, tubes were filled with 1 ml of PBS; 80 μg/ml dispersin B solution; or 0.1%, 0.5%, 1% or 2.5% BPO, as indicated. After 15 min at 37°C, tubes were rinsed three times with 3 ml of saline and filled with 1 ml of saline. Biofilm bacteria were detached from the walls of the tubes by sonication for 30 sec using an IKA Labortechnik sonicator set to 50% power and 50% duty cycle. Control experiments showed that this sonication treatment did not affect the viability of *C. acnes* or *S. epidermidis* cells (data not shown). Sonicates were serially diluted in saline. To enumerate *S. epidermidis*, dilutions were plated on Tryptic Soy agar and incubated in air. To enumerate *C. acnes*, dilutions were plated on Tryptic Soy agar supplemented with 20 μg/ml erythromycin and incubated anaerobically.

### Crystal violet binding assay

Biofilms were rinsed vigorously with tap water and stained for 1 min with 1 ml of Gram’s crystal violet. Tubes were then rinsed with tap water to remove the unbound dye and airdried. To quantitate crystal violet binding, tubes were filled with 1 ml of 33% acetic acid, incubated at room temperature for 30 min, and mixed by vortex agitation. A volume of 200 μl of the dissolved dye was transferred to the well of a 96-well microtiter plate and its absorbance at 620 nm was measured in an automated microplate reader. Tubes containing sterile broth were incubated and processed along with the inoculated tubes to serve as controls. The amount of biofilm detachment was quantitated using the formula 1 - (*A*620_Dispersin B_/*A*620_No enzyme_) × 100.

### Statistics and reproducibility of results

All experiments were performed in duplicate or triplicate tubes. All experiments were performed on at least three occasions with similar results. The significance of differences between means was calculated using a Student’s *t*-test. A *P-*value < 0.01 was considered significant.

## RESULTS

### Detachment of *S. epidermidis*/*C. acnes* dual-species biofilms by dispersin B solution and dispersin B hydrogel

To confirm that dispersin B solution and dispersin B hydrogel cause detachment of *S. epidermidis*/*C. acnes* dual-species biofilms grown in glass tubes, biofilms were treated with a 80 μg/ml dispersin B solution, or a hydrogel containing 80 μg/ml dispersin B, for 15 min. The amount of biofilm biomass remaining after treatment was quantitated using a crystal violet binding assay (Fig. 1). Both the dispersin B solution and dispersin B hydrogel treatments caused a significant reduction in biofilm biomass (94% and 78%, respectively). These results are consistent with those of a previous study showing that a 100 μg/ml solution of dispersin B efficiently detached aerobic *S. epidermidis*/*C. acnes* dual-species biofilms from glass tubes (Kaplan, 2023b). The biofilm detaching activity of dispersin B hydrogel was evidently due to the enzyme and not the other components of the hydrogel because a blank hydrogel (no enzyme) did not cause significant detachment of *S. epidermidis*/*C. acnes* dual-species biofilms as evidenced by visual inspection of the biofilms after treatment (Fig. 2).

**Figure 1.**
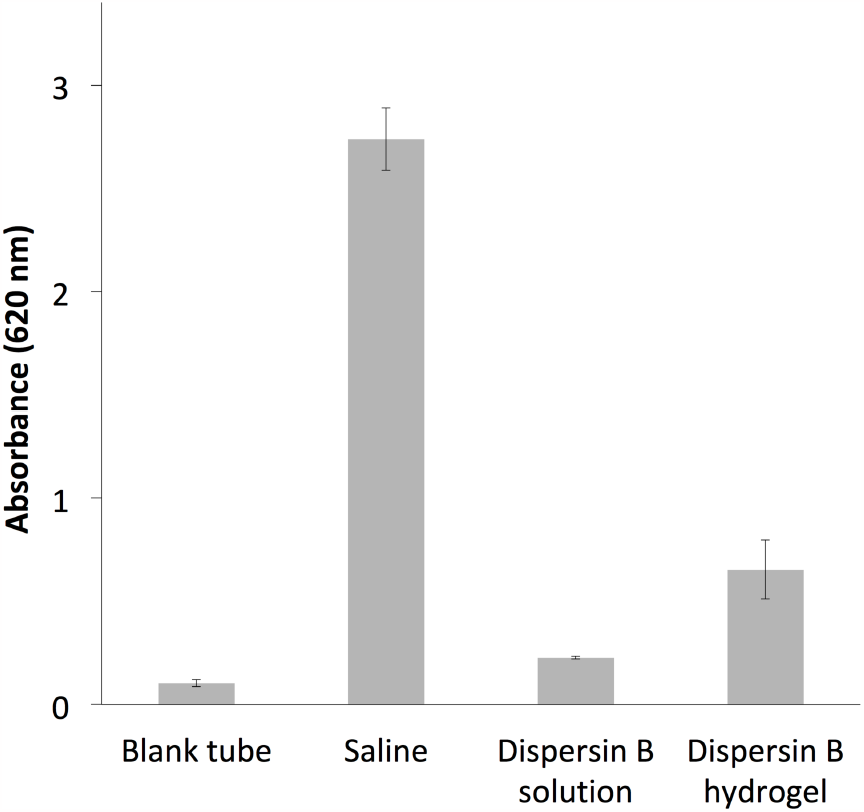
**Detachment of *S. epidermidis*/*C. acnes* dual-species biofilms by dispersin B solution and dispersin B hydrogel. Both agents contained 80 μg/ml of enzyme. Biofilms were rinsed and treated with the indicated agent for 15 min, then stained with crystal violet. The amount of bound dye (Absorbance at 620 nm) is proportional to the amount of biofilm biomass. Control tubes were treated with PBS (Saline). Values show mean and range for triplicate tubes for each condition**.

**Figure 2.**
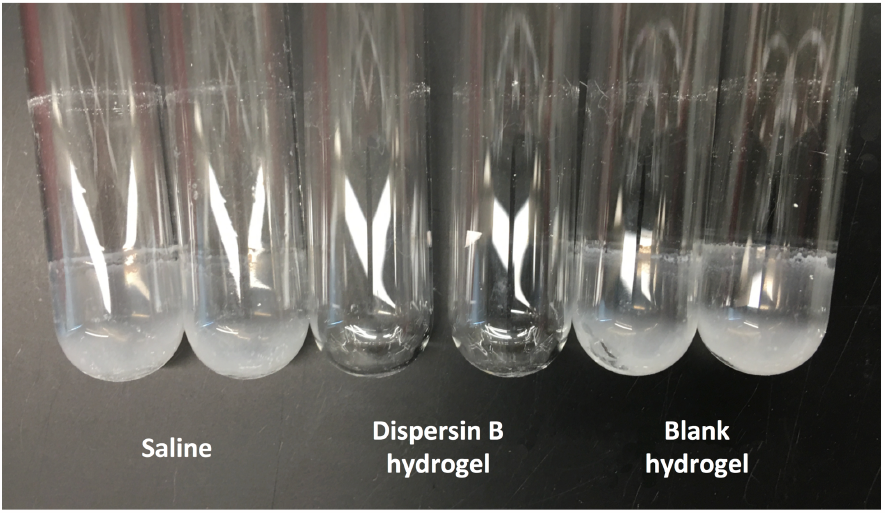
**Detachment of *S. epidermidis*/*C. acnes* dual-species biofilms by dispersin B hydrogel. Biofilms were rinsed and treated with the indicated agent for 15 min, then rinsed and photographed. Control tubes were treated with PBS (Saline). Duplicate tubes for each condition are shown**.

### Pre-treatment of *S. epidermidis*/*C. acnes* dual-species biofilms with dispersin B solution renders them susceptible to benzoyl peroxide killing

*S. epidermidis*/*C. acnes* dual-species biofilms were subjected to a series of two-step treatments consisting of PBS, 80 μg/ml dispersin B solution, or 1% benzoyl peroxide in different combinations. After treatment, biofilms were detached from the walls of the tube by sonication and *S. epidermidis* and *C. acnes* CFUs were enumerated independently (Fig. 3). Dispersin B treatment alone caused a >1 log reduction in *S. epidermidis* CFUs and a >2 log reduction in *C. acnes* CFUs. Treatment of biofilms with 1% benzoyl peroxide alone caused no reduction in *S. epidermidis* CFUs and a 1 log reduction in *C. acnes* CFUs. However, treatment of biofilms with dispersin B followed by benzoyl peroxide caused a > 2 log reduction in *S. epidermidis* CFUs and a > 5 log reduction in *C. acnes* CFUs. These results suggest that dispersin B sensitizes both *S. epidermidis* and *C. acnes* to benzoyl peroxide killing when cultured aerobically in a dual-species biofilm.

**FIG. 3.**
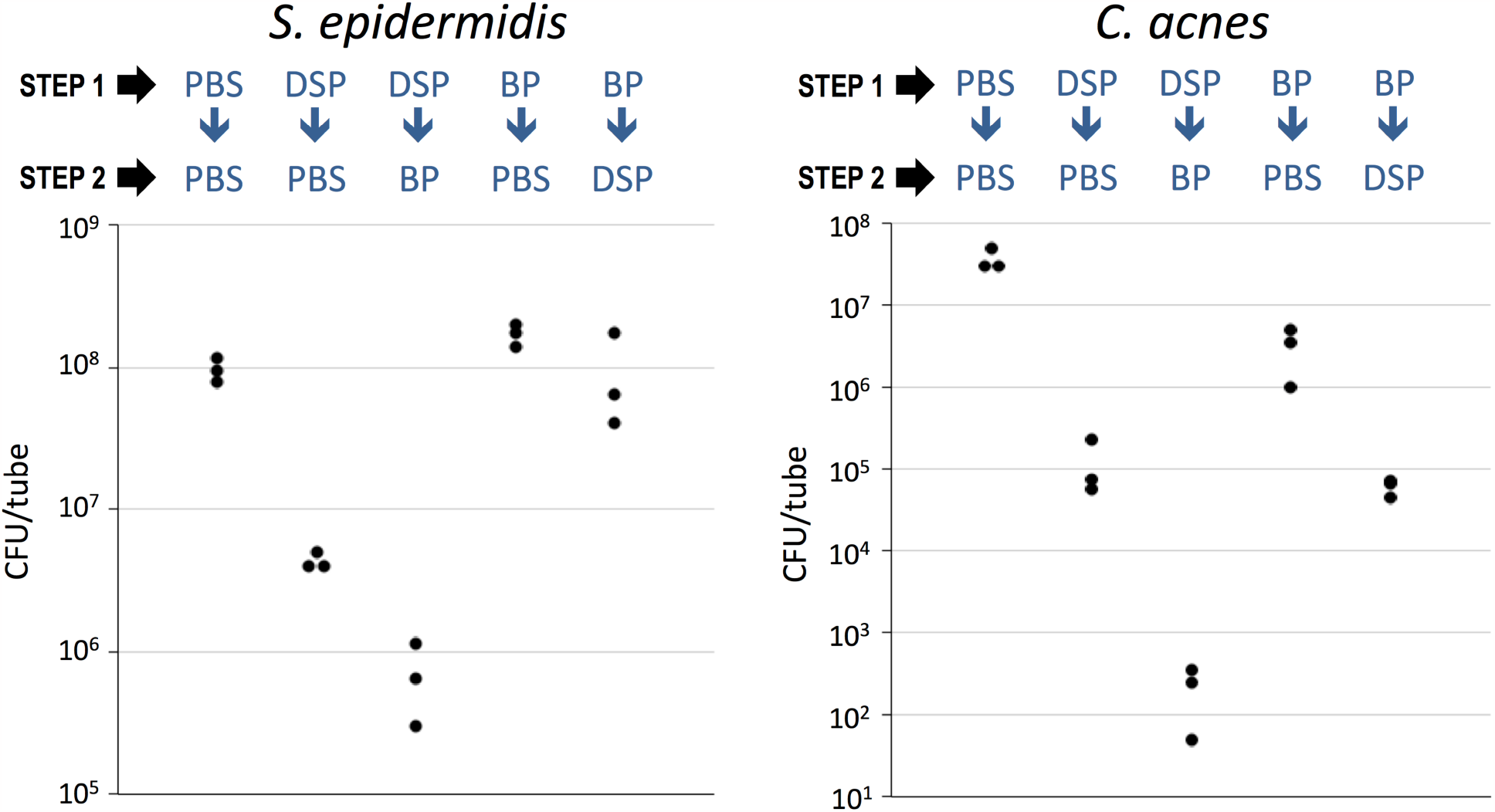
**Pre-treatment of *S. epidermidis*/*C. acnes* dual-species biofilms with dispersin B solution renders them sensitive to killing by benzoyl peroxide. 24-h-old aerobic *S. epidermidis*/*C. acnes* dual-species biofilms were rinsed and treated with phosphate buffered saline (PBS), 80 μg/ml dispersin B solution (DSP), or 1% benzoyl peroxide (BP). After 15 min, biofilms were rinsed and subjected to a second 15-min treatment with PBS, DSP or BP in the indicated combinations. Graphs show CFU/tube values independently for *S. epidermidis* (left panel) and *C. acnes* (right panel) from triplicate tubes for each treatment. Each dot represents one tube**.

Treatment of biofilms with benzoyl peroxide followed by dispersin B resulted in no significant detachment or killing of *S. epidermidis*, suggesting that residual benzoyl peroxide may partially inactivate or inhibit dispersin B. Benzoyl peroxide treatment followed by dispersin B treatment resulted in a 3 log reduction in *C. acnes* CFUs compared to a > 5 log reduction when biofilms were treated with dispersin B followed by benzoyl peroxide. In a similar experiment, dispersin B treatment followed by 1% benzoyl peroxide caused significantly more *C. acnes* detachment and killing than 2.5% benzoyl peroxide alone (Fig. 4), suggesting that dispersin B treatment may lower the minimum effective dose of benzoyl peroxide against *C. acnes*.

**FIG. 4.**
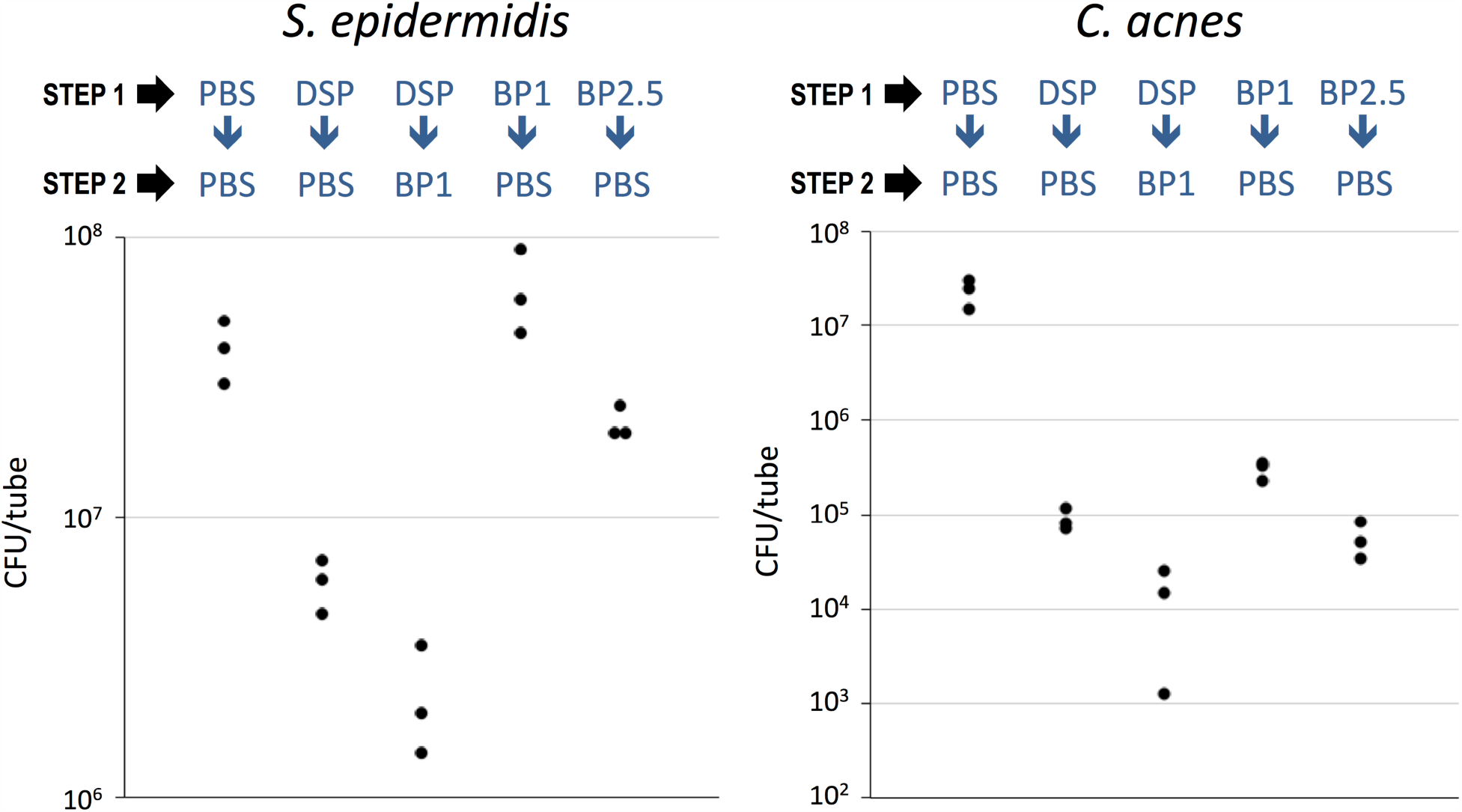
**Dispersin B solution + 1% benzoyl peroxide kills *C. acnes* biofilms as efficiently as 2**.**5% benzoyl peroxide alone. 24-h-old aerobic *S. epidermidis*/*C. acnes* dual-species biofilms were rinsed and treated with phosphate buffered saline (PBS), 80 μg/ml dispersin B solution (DSP), or 1% or 2**.**5% benzoyl peroxide (BP1 and BP2**.**5, respectively). After 15 min, biofilms were rinsed and subjected to a second 15-min treatment with PBS or BP1 in the indicated combinations. Graphs show CFU/tube values independently for *S. epidermidis* (left panel) and *C. acnes* (right panel) from triplicate tubes for each treatment. Each dot represents one tube**.

### Dispersin B hydrogel acts synergistically with benzoyl peroxide to kill *C. acnes* in *S. epidermidis*/*C. acnes* dual-species biofilms

To determine whether dispersin B hydrogel can sensitize *S. epidermidis*/*C. acnes* dual-species biofilms to benzoyl peroxide killing, biofilms were subjected to a two-step treatment where the first step consisted of PBS, or dispersin B hydrogel containing 40 or 80 μg/ml dispersin B, and the second step consisted of PBS, 1% benzoyl peroxide, or 2.5% benzoyl peroxide. After treatment, biofilms were detached from the walls of the tube by sonication and *S. epidermidis* and *C. acnes* CFUs were enumerated independently (Fig. 5). Dispersin B hydrogel alone at 40 or 80 μg/ml resulted in a > 1 log reduction in *S. epidermidis* CFUs, while benzoyl peroxide alone at 1% or 2.5% exhibited no significant killing of *S. epidermidis* biofilms (Fig. 5, left panel). Dispersin B hydrogel at 40 or 80 μg/ml did not sensitize *S. epidermidis* to 1% or 2.5% benzoyl peroxide killing under these conditions (Fig. 5, left panel). In contrast, dispersin B hydrogel alone at 40 or 80 μg/ml resulted in a > 2 log reduction in *C. acnes* CFUs, benzoyl peroxide alone at 1% or 2.5% caused > 1 log killing of *C. acnes* biofilms, and dispersin B hydrogel at 40 or 80 μg/ml followed by benzoyl peroxide at 1% or 2.5% caused > 6 log reduction in *C. acnes* CFUs (Fig. 5, right panel). These results suggest that dispersin B hydrogel acts synergistically with benzoyl peroxide to kill *C. acnes* in *S. epidermidis*/*C. acnes* dual-species biofilms. To determine the lowest concentrations of dispersin B and benzoyl peroxide needed to eradicate *C. acnes* from *S. epidermidis*/*C. acnes* dual-species biofilms, biofilms were treated with dispersin B hydrogel containing 20, 10 or 5 μg/ml dispersin B, followed by benzoyl peroxide at 0.5% or 0.1% (Fig. 6). Dispersin B hydrogel alone at enzyme concentrations of 20, 10 or 5 μg/ml resulted in a > 1 log reduction in *S. epidermidis* CFUs and a 2 log reduction in *C. acnes* CFUs (Fig. 6), while benzoyl peroxide alone at 0.5% or 0.1% exhibited no significant killing of *S. epidermidis* biofilms (Fig. 6, left panel) and a 1-2 log reduction in *C. acnes* CFUs (Fig. 6, right panel). Dispersin B hydrogel at 20, 10 or 5 μg/ml of enzyme did not sensitize *S. epidermidis* to killing by 0.5% or 0.1% benzoyl peroxide (Fig. 6, left panel). In contrast, treatment of biofilms with dispersin B hydrogel at enzyme concentrations 20, 10 or 5 μg/ml resulted in a > 3 log increase in *C. acnes* killing by both 0.5% and 0.1% benzoyl peroxide compared to 0.5% and 0.1% benzoyl peroxide alone (Fig. 6, right panel).

**FIG. 5.**
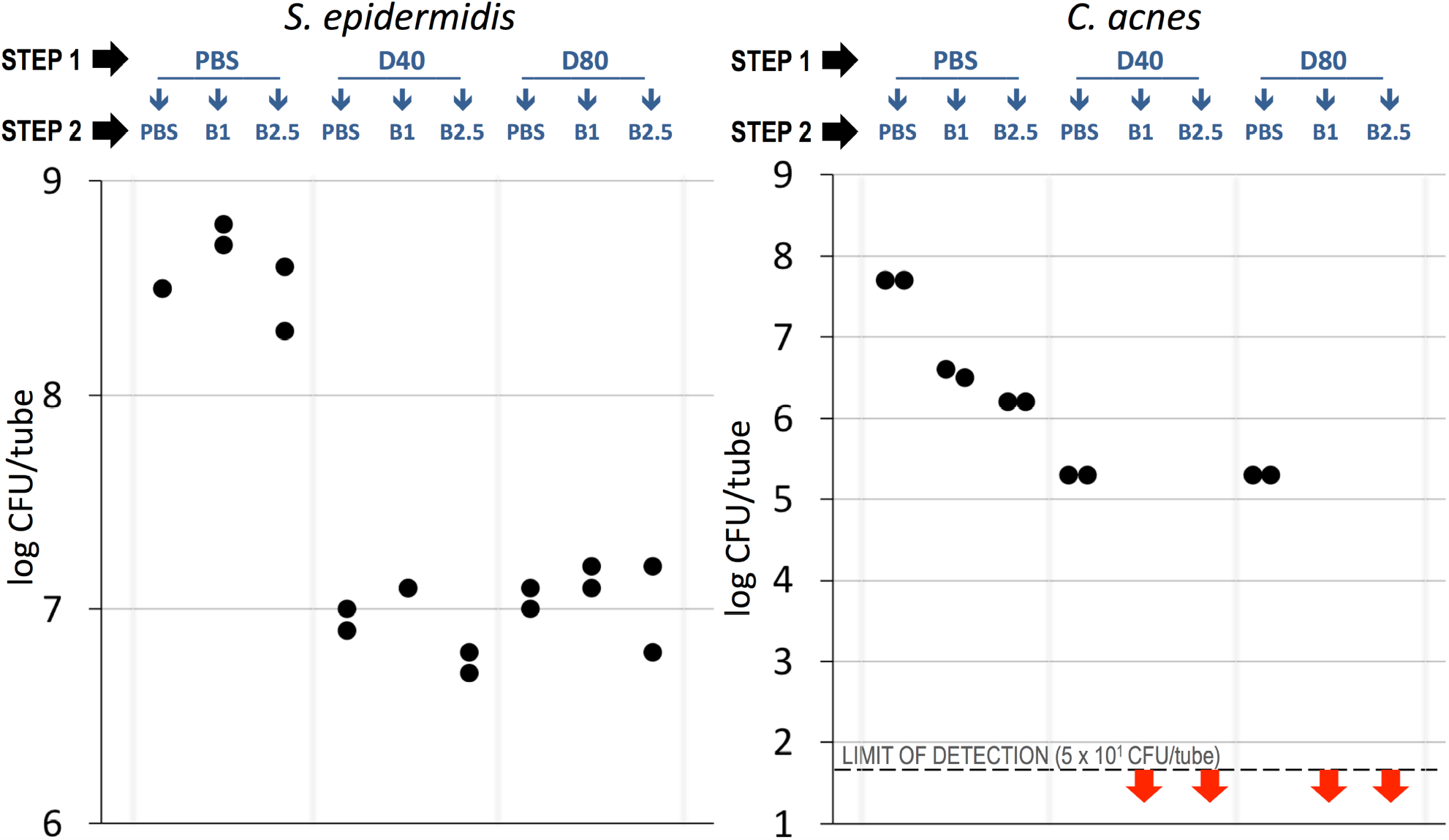
**Dispersin B hydrogel sensitizes *C. acnes* biofilms to benzoyl peroxide killing. 24-h-old aerobic *S. epidermidis*/*C. acnes* dual-species biofilms were rinsed and treated with phosphate buffered saline (PBS) or dispersin B hydrogel containing 40 or 80 μg/ml dispersin B (D40 and D80, respectively). After 15 min, biofilms were rinsed and subjected to a second 15-min treatment with PBS or benzoyl peroxide at 1% or 2**.**5% (BP1 and BP2**.**5, respectively), in the combinations shown. Graphs show CFU/tube values independently for *S. epidermidis* (left panel) and *C. acnes* (right panel) from duplicate tubes for each treatment. Red arrows indicate that the CFU values were below the limit of detection (50 CFUs). Each dot represents one tube**.

**FIG. 6.**
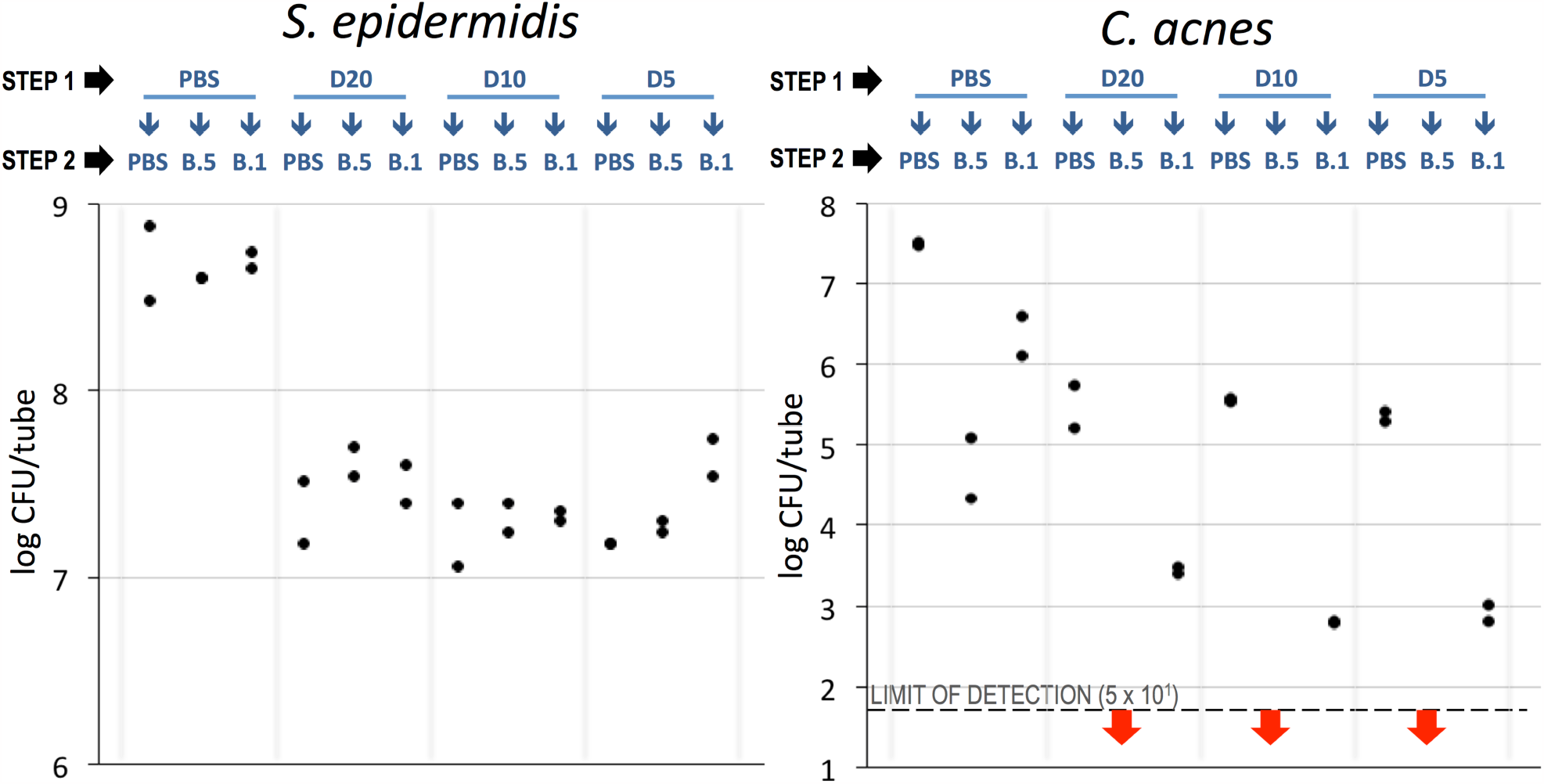
**Dispersin B hydrogel sensitizes *C. acnes* biofilms to benzoyl peroxide killing. 24-h-old aerobic *S. epidermidis*/*C. acnes* dual-species biofilms were rinsed and treated with phosphate buffered saline (PBS) or dispersin B hydrogel containing 20, 10 or 5 μg/ml dispersin B (D20, D10, and D5 respectively). After 15 min, biofilms were rinsed and treated with PBS or benzoyl peroxide at 0**.**5% or 0**.**1% (BP**.**5 and BP**.**1, respectively). Graphs show CFU/tube values independently for *S. epidermidis* (left panel) and *C. acnes* (right panel) from duplicate tubes for each treatment. Each dot represents one tube**.

## DISCUSSION

Benzoyl peroxide (BPO) has been a mainstay of acne treatment since the 1950s. BPO is an oxidizing agent that non-specifically oxidizes bacterial surface proteins, resulting in a loss of protein structure and rapid cell death. BPO exhibits bactericidal activity against both *C. acnes* and *S. epidermidis* (Cove & Holland, 1983). The advantages of BPO are that BPO circumvents the use of antibiotics, it is effective against antibiotic-resistant strains, and resistance to BPO has not been reported. A major disadvantage of BPO, however, is its skin irritability, which causes some acne patients to switch to lower doses that are less effective because they require a longer contact time to kill *C. acnes* (Boonchaya *et al*., 2022).

Previous studies showed that dispersin B sensitized anaerobic *C. acnes* mono-species biofilms to killing by benzoyl peroxide and tetracycline (Kaplan, 2023a) and dispersed aerobic *C. acnes*/*S. epidermidis* dual-species biofilms (Kaplan, 2023b). In the present study we showed that dispersin B sensitizes aerobic *C. acnes*/*S. epidermidis* dual-species biofilms to benzoyl peroxide killing. This activity is consistent with the biocide sensitization activities exhibited by dispersin B against other species of bacteria (Ganeshnarayan *et al*., 2009; Itoh *et al*., 2005; Izano *et al*., 2007; Parise *et al*., 2007). Because of the high specificity of dispersin B for PNAG (Eddenden & Nitz, 2022), these findings suggest that PNAG plays an important role in *C. acnes* benzoyl peroxide resistance. Dispersin B may be a useful adjunct to BPO for the treatment and prevention of acne because it may decrease the minimum effective dose of BPO and thereby minimize skin irritation.

## ACKNOWLEDGEMENTS

The author thanks BEI Resources (Manassas, Virginia, USA) for providing *C. acnes* strain HL086PA1; Kane Biotech Inc. (Winnipeg, Manitoba, Canada) for providing dispersin B solution, dispersin B hydrogel, and blank hydrogel; and Katie DeCicco-Skinner (American University) for continued support.

## REFERENCES

Achermann, Y., E. J. Goldstein, T. Coenye, and M. E. Shirtliff. 2014. Propionibacterium acnes: from commensal to opportunistic biofilm-associated implant pathogen. Clin Microbiol Rev 27 (3): 419–40. 10.1128/CMR.00092-13.

Allen, H. B., and J. L. Mueller. 2011. A novel finding in atopic dermatitis: film-producing Staphylococcus epidermidis as an etiology. Int J Dermatol 50 (8): 992–3. doi: 10.1111/j.1365-4632.2010.04648.x.

Barnard, E., B. Shi, D. Kang, N. Craft, and H. Li. 2016. The balance of metagenomic elements shapes the skin microbiome in acne and health. Sci Rep 6: 39491. 10.1038/srep39491. https://www.ncbi.nlm.nih.gov/pubmed/28000755.

Bayston, R., W. Ashraf, R. Barker-Davies, E. Tucker, R. Clement, J. Clayton, B. J. Freeman, and B. Nuradeen. 2006. Biofilm formation by Propionibacterium acnes on biomaterials in vitro and in vivo: impact on diagnosis and treatment. J Biomed Mater Res A 81 (3): 705–9. 10.1002/jbm.a.31145.

Bek-Thomsen, M., H. B. Lomholt, and M. Kilian. 2008. Acne is not associated with yet-uncultured bacteria. J Clin Microbiol 46 (10): 3355–60. 10.1128/JCM.00799-08.

Boonchaya, P., S. Rojhirunsakool, N. Kamanamool, S. Khunkhet, S. Yooyongsatit, M. Udompataikul, and M. Taweechotipatr. 2022. Minimum contact time of 1.25%, 2.5%, 5%, and 10% benzoyl peroxide for a bactericidal effect against Cutibacterium acnes. Clin Cometic Invest Dermatol 15: 403–409. doi: 10.2147/CCID.S359055.

Bütthner, H., D. Mack, and H. Rohde. 2015. Structural basis of Staphylococcus epidermidis biofilm formation: mechanisms and molecular interactions. Front Cell Infect Microbiol 5: 14. 10.3389/fcimb.2015.00014.

Chaignon, P., I. Sadovskaya, C. Ragunath, N. Ramasubbu, J. B. Kaplan, and S. Jabbouri. 2007. Susceptibility of staphylococcal biofilms to enzymatic treatments depends on their chemical composition. Appl Microbiol Biotechnol 75: 125–132. doi: 10.1007/s00253-006-0790-y.

Chen, K. -M., M. -K. Chiang, M. Wang, H. -C. Ho, M. -C. Lu, Y. -C. Lai. 2014. The role of pgaC in Klebsiella pneumoniae virulence and biofilm formation. Miccrob Pathog 77: 89–99. doi: 10.1016/j.micpath.2014.11.005.

Claudel, J. P., N. Auffret, M. T. Leccia, F. Poli, S. Corvec, and B. Dréno. 2019. Staphylococcus epidermidis: a potential new player in the physiopathology of acne? Dermatology 235 (4): 287–294. 10.1159/000499858.

Costerton, J. W., P. S. Stewart, and E. P. Greenberg. 1999. Bacterial biofilms: a common cause of persistent infection. Sceince 284 (5418): 1318–22. doi: 10.1126/science.284.5418.1318.

Cove, J. H., and K. T. Holland. 1983. The effect of benzoyl peroxide on cutaneous micro-organisms in vitro. J Apl Bacteriol 54 (3): 379–82. 10.1111/j.1365-2672.1983.tb02631.x.

Cywes-Bentley, C., D. Skurnik, T. Zaidi, D. Roux, R. B. Deoliveira, W. S. Garret, X. Lu, J. O’Malley, K. Kinzel, A. Rey, C. Perrin, R. N. Fichorova, A. K. Kayatani, T. Maira-Litràn, M. L. Gening, Y. E. Tsvetkov, N. E. Nifantiev, L. O. Bakaletz, S. I. Pelton, D. T. Golenbock, and G. B. Pier. 2013. Antibody to a conserved antigenic target is protective against diverse prokaryotic and eukaryotic pathogens. Proc Natl Acad Sci U S A 110 (24): E2209–18. 10.1073/pnas.1303573110.

Dréno, B. R. Martin, D. Moyal J. B. Henley, A. Khammari, and S. Seité. 2017. Skin microbiome and acne vulgaris: Staphylococcus, a new actor in acne. Exp Dermatol 88 (26): 798–803. 10.1111/exd.13296.

Eddenden, A., and M. Nitz. 2022. Applications of an inactive Dispersin B probe to monitor biofilm polysaccharide production. Methods Enzymol 665:209–231. doi: 10.1016/bs.mie.2021.11.006.

Evans, S. M., A.E. Schrlau, A. A. Chalian, P. Zhang, and C. J. Koch. 2006. Oxygen levels in normal and previously irradiated human skin as assessed by EF5 binding. J Invest Dermatol 126(12): 2596–2606. doi: 10.1038/sj.jid.5700451.

Fitz-Gibbon, S., S. Tomida, B. -H. Chiu, L. Nguyen, C. Du, M. Liu, D. Elashoff, M. C. Erfe, A. Loncaric, J. Kim, R. L. Modlin, J. F. Miller, E. Sodergren, N. Craft, G. M. Weinstock, and H. Li. 2013. Propionibacterium acnes strain populations in the human skin microbiome associated with acne. J Invest Dermatol 133(9): 2152–2160. doi: 10.1038/jid.2013.21.

Ganeshnarayan, K., S. M. Shah, M. R. Libera, A. Santostefano, and J. B. Kaplan. 2009. Poly-N-acetylglucosamine matrix polysaccharide impedes fluid convection and transport of the cationic surfactant cetylpyridinium chloride through bacterial biofilms. Appl Environ Microbiol 75 (5): 1308–14. 10.1128/AEM.01900-08.

Greydanus, D. E., R. Azmeh, M. D. Cabral, C. A. Dickson, and D. R. Patel. 2021. Acne in the first three decades of life: an update of a disorder with profound implications for all decades of life. Dis Mon 67 (4): 101103. 10.1016/j.disamonth.2020.101103.

Itoh, Y., X. Wang, B. J. Hinnebusch, J. F. Preston, III, and T. Romeo. 2005. Depolymerization of β-1,6-N-acetyl-D-glucosamine disrupts the integrity of diverse bacterial biofilms. J Bacteriol 187 (1): 382–7. doi: 10.1128/JB.187.1.382-387.2005

Izano, E. A., I. Sadovskaya, E. Vinogradov, M. H. Mulks, K. Velliyagounder, C. Ragunath, W. B. Kher, N. Ramasubbu, S. Jabbouri, M. B. Perry, and J. B. Kaplan. 2007. Poly-N-acetylglucosamine mediates biofilm formation and antibiotic resistance in Actinobacillus pleuropneumoniae. Microb Pathog 43 (1): 1–9. 10.1016/j.micpath.2007.02.004.

Jahns, A. C., B. Lundskog, R. Ganceviciene, R. H. Palmer, I. Golovleva, C. C. Zouboulis, A. McDowell, S. Patrick, and O. A. Alexeyev. 2012. An increased incidence of Propionibacterium acnes biofilms in acne vulgaris: a casecontrol study. Br J Dermatol 167 (1): 50–8. 10.1111/j.1365-2133.2012.10897.x.

Kaplan, J. B. 2023a. Poly-N-acetylglucosamine mediates Cutibacterium acnes biofilm formation and biocide resistance. bioRxiv. doi: 10.1101/2023.10.10.558046.

Kaplan, J. B. 2023b. Staphylococcus epidermidis enables Cutibacterium acnes to form biofilms under aerobic conditions. bioRxiv. doi: 10.1101/2023.10.12.562081.

Kaplan, J. B., K. D. Mlynek, H. Hettiarachchi, Y. A. Alamneh, L. Biggemann, D. V. Zurawski, C. C. Black, C. E. Bane, R. K. Kim, and M. S. Granick. 2018. Extracellular polymeric substance (EPS)-degrading enzymes reduce staphylococcal surface attachment and biocide resistance on pig skin in vivo. PloS ONE 13(10): e0205526. 10.1371/journal.pone.0205526.

Kaplan, J. B., C. Ragunath, K. Velliyagounder, D. H. Fine, andN. Ramasubbu. 2004. Enzymatic detachment of Staphylococcus epidermidis biofilms. Antmicrob Agents Chemother 48(7): 2633–2636. 10.1128/aac.48.7.2633-2636.2004.

Kropec, A., T. Maira-Litran, K. K. Jefferson, M. Grout, S. E. Cramton, F. Götz, D. A. Goldmann, and G. B. Pier. 2005. Poly-N-acetylglucosamine production in Staphylococcus aureus is essential for virulence in murine models of systemic infection. Infect Immun 73 (10): 6868–76. 10.1128/IAI.73.10.6868-6876.2005.

Lee, Y. B., E. J. Byun, and H. S. Kim. 2019. Potential role of the microbiome in acne: a comprehensive review. J Clin Med 8 (7). 10.3390/jcm8070987.

Lee, J. -H., J. B. Kaplan, W. Y. Lee. 2008. Microfluidic devices for studying growth and detachment of Staphylococcus epidermidis biofilms. Biomed Microdevices 10(4): 489–98. doi: 10.1007/s10544-007-9157-0.

Mayslich, C., P. A. Grange, and N. Dupin. 2021. Cu8bacterium acnes as an opportunistic pathogen: an update of its virulence-associated factors. Microorganisms 9 (2). 10.3390/microorganisms9020303.

McLaughlin, J., S. Watterson, A. M. Layton, A. J. Bjourson, E. Barnard, and A. McDowell. 2019. Propionibacterium acnes and acne vulgaris: new insights from the integration of population genetic, multi -omic, biochemical and host-microbe studies. Microorganisms 7 (5). 10.3390/microorganisms7050128.

O’Neill, A. M., and R. L. Gallo. 2018. Host-microbiome interactions and recent progress into understanding the biology of acne vulgaris. Microbiome 6 (1): 177. 10.1186/s40168-018-0558-5.

Oho, M. 2009. Staphylococcus epidermidis – the “accidental” pathogen. Nat Rev Microbiol 7(8): 555–567. doi: 10.1038/nrmicro2182.

Parise, G., M. Mishra, Y. Itoh, T. Romeo, and R. Deora. 2007. Role of a putative polysaccharide locus in Bordetella biofilm development. J Bacteriol 189 (3): 750–60. 10.1128/JB.00953-06.

Shanmugam, M., P. Gopal, F. El Abbar, H. C. Schreiner, J. B. Kaplan, D. H. Fine, and N. Ramasubbu. 2015. Role of exopolysaccharide in Aggregatibacter actinomycetemcomitans-induced bone resorption in a rat model for periodontal disease. PloS ONE 10:e117487. doi: 10.1371/journal.pone.0117487.

Soliman, C., A. K. Walduck, E. Yuriev, J. S. Richards, C. Cywes-Bentley, G. B. Pier, and P. A. Ramsland. 2018. Structural basis for antibody targeting of the broadly expressed microbial polysaccharide poly-N-acetylglucosamine. J Biol Chem 293 (14): 5079–5089. 10.1074/jbc.RA117.001170.

Subashchandrabose, S. R. M. Leveque, R. N.Kirkwood, M. Kiupel, and M. H. Mulks. 2013. The RNA chaperone Hfq promotes fitness of Actinobacillus pleuropneumoniae during porcine pleuropneumonia. Infect Immun 81 (8): 2952–61. 10.1128/IAI.00392-13.

